# A partnership of the lipid scramblase XK and of the lipid transfer protein VPS13A at the plasma membrane

**DOI:** 10.1101/2022.03.30.486314

**Authors:** Andrés Guillén Samander, Yumei Wu, S. Sebastian Pineda, Francisco J. García, Julia N. Eisen, Marianna Leonzino, Berrak Uğur, Manolis Kellis, Myriam Heiman, Pietro De Camilli

## Abstract

Chorea-acanthocytosis and McLeod syndrome are diseases with shared clinical manifestations caused by mutations in VPS13A and XK, respectively. Key features of these conditions are the degeneration of caudate neurons and the presence of abnormally shaped erythrocytes. XK belongs to a family of plasma membrane (PM) lipid scramblases whose action results in exposure of PtdSer at the cell surface. VPS13A is an ER-anchored lipid transfer protein with a putative role in the transport of lipids at contacts of the ER with other membranes. Recently VPS13A and XK were reported to interact by still unknown mechanisms. So far, however, there is no evidence for a colocalization of the two proteins at contacts of the ER with the PM, where XK resides, as VPS13A was shown to be localized at contacts between the ER and either mitochondria or lipid droplets. Here we show that VPS13A can also localize at ER-PM contacts via the binding of its PH domain to a cytosolic loop of XK, that such interaction is regulated by an intramolecular interaction within XK and that both VPS13A and XK are highly expressed in the caudate neurons. Binding of the PH domain of VPS13A to XK is competitive with its binding to intracellular membranes that mediate other tethering functions of VPS13A. Our findings support a model according to which VPS13A-dependent lipid transfer between the ER and the PM is coupled to lipid scrambling within the PM. They raise the possibility that defective cell surface exposure of PtdSer may be responsible for neurodegeneration.

## Introduction

Chorea-acanthocytosis (ChAc) and McLeod syndrome (MLS) are two similar diseases characterized by progressive degeneration of the caudate nucleus leading to chorea and other movement defects, and by abnormally shaped erythrocytes (acanthocytes). Chorea acanthocytosis is due to loss-of-function of VPS13A(1, 2), the founding member of a family of lipid transport proteins, also called chorein motif protein family. Members of this family, which also comprise the autophagy factor ATG2, are localized at membrane contact sites where they are thought to transfer lipids unidirectionally between adjacent bilayers of different organelles by a bridge-like mechanism(3, 4). McLeod syndrome is due instead to loss-of-function of XK(5), a member of a family of lipid scramblases whose function is to collapse the heterogeneity of the lipid composition of the two leaflets of the plasma membrane (PM)(6). A consequence of this scrambling is the exposure to the cell surface of PtdSer, a phospholipid that is recognized by phagocytic cells and which is normally concentrated in the cytosolic leaflet of the PM by the action of flippases(7).

Consistent with the similarity of the clinical conditions resulting from mutations in VPS13A and XK, there is evidence suggesting that the two proteins are functional partners(8–10). Thus, it is of great interest that both proteins are implicated in lipid transport. More importantly, as chorein motif proteins do not penetrate lipid bilayers but are thought to achieve net transfer of lipids between cytosolic leaflets of the donor and receiving membranes, they are expected to require the cooperation of scramblases to allow equilibration of lipid mass between the bilayer leaflets. Accordingly, there is evidence for the partnership of the chorein motif protein ATG2 with lipid scramblases in the growth of the autophagosome membrane(4, 11, 12). A cooperation between VPS13A and XK, which has been recently confirmed to have scramblase activity by in vitro studies(13), would represent another example of such partnership, providing clues to mechanisms of disease in chorea-acanthocytosis and McLeod syndrome. Strong support for this possibility came by the demonstration of an interaction between these two proteins: i) studies of McLeod syndrome erythrocytes revealed that lack of XK results in a loss of VPS13A in their membranes(8), as expected if VPS13 and XK are part of a same complex; ii) their interaction was supported by biochemical experiments(8–10); iii) the localization of overexpressed VPS13A at contacts between ER and lipid droplets was shown to abolished by co-overexpression of XK resulting in a localization VPS13A along with XK throughout the ER(9).

So far, however, there has been no evidence from imaging studies in mammalian cells that VPS13A can colocalized with XK at the plasma membrane, where XK is expected to act. Studies in HeLa cells with tagged-VPS13A - both overexpressed VPS13A(9, 14–16), and VPS13A expressed at endogenous levels(14)-have shown that this protein is localized at contacts between the ER and mitochondria and also at contacts between the ER and lipid droplets when these organelles are present. These studies have further shown that this localization is mediated by the interaction of the N-terminal region of VPS13A with the MSP domain of the ER protein VAP and of its C-terminal region with a yet to be defined binding site on mitochondria and on lipid droplets(14, 16). An interaction of VPS13A with mitochondria was also supported by the identification of this protein as a hit in a screen for mitochondria neighbors by proximity biotinylation(17, 18). Based on these findings, it has been proposed that VPS13A, like its paralog VPS13D, mediates transport of lipids to mitochondria which are organelles not connected by vesicular transport to the ER, where most membrane lipids are synthesized(14, 19).

Goal of this study was to determine how VPS13A interacts with XK and to determine whether it can be localized with XK at the plasma membrane.

## Results

### Predicted structure of VPS13A

As a premise to the elucidation of how VPS13A interacts with XK, we capitalized on Alphafold2-based algorithms(20) to complement previous structural studies of VPS13 family proteins based on crystallography(14), negative staining EM(21) and cryoEM(22) and to predict VPS13A full-length structure. Based on such prediction, VPS13A has the shape of a ∼22nm rod whose core is represented by an elongated twisted *β*-sheet surrounded at its C-terminal end by other folded modules (Fig. 1A-C and Movie S1). The rod harbors a groove that runs along its entire length and has a hydrophobic floor, supporting the hypothesis that VPS13A (like other VPS13 proteins), bridges two lipid bilayers to allow bulk flow of lipids between them(3, 4, 23, 24). VPS13A is anchored to the ER via a VAP binding FFAT motif(14, 16, 25) found at about ∼6nm from the N-terminal end of the rod, which comprises the so-called chorein motif. As in other VPS13 isoforms, the modules that surround the C-terminal region of VPS13A are the arc-like VPS13 Adaptor Binding (VAB), a bundle of *α*-helices (called ATG2-C domain) that have similarity to a similar bundle found in ATG2-C(26), and a PH domain(27). An additional previously unnoticed small folded module is the WWE domain, which is also present in VPS13C(23), the closest human paralogue of VPS13A and in the metazoan orthologs of VPS13A/C. WWE domains in other proteins were shown to bind Poly-ADP-Ribose (PAR)(28). Both the VAB and the WWE domains are outpocketing of the elongated *β*-sheet rod, as the region directly downstream to these domains (orange in Fig. 1A and B), including the so-called Apt-1 domain(29), represents the C-terminal portion of the *β*-sheet rod.

**Figure 1.**
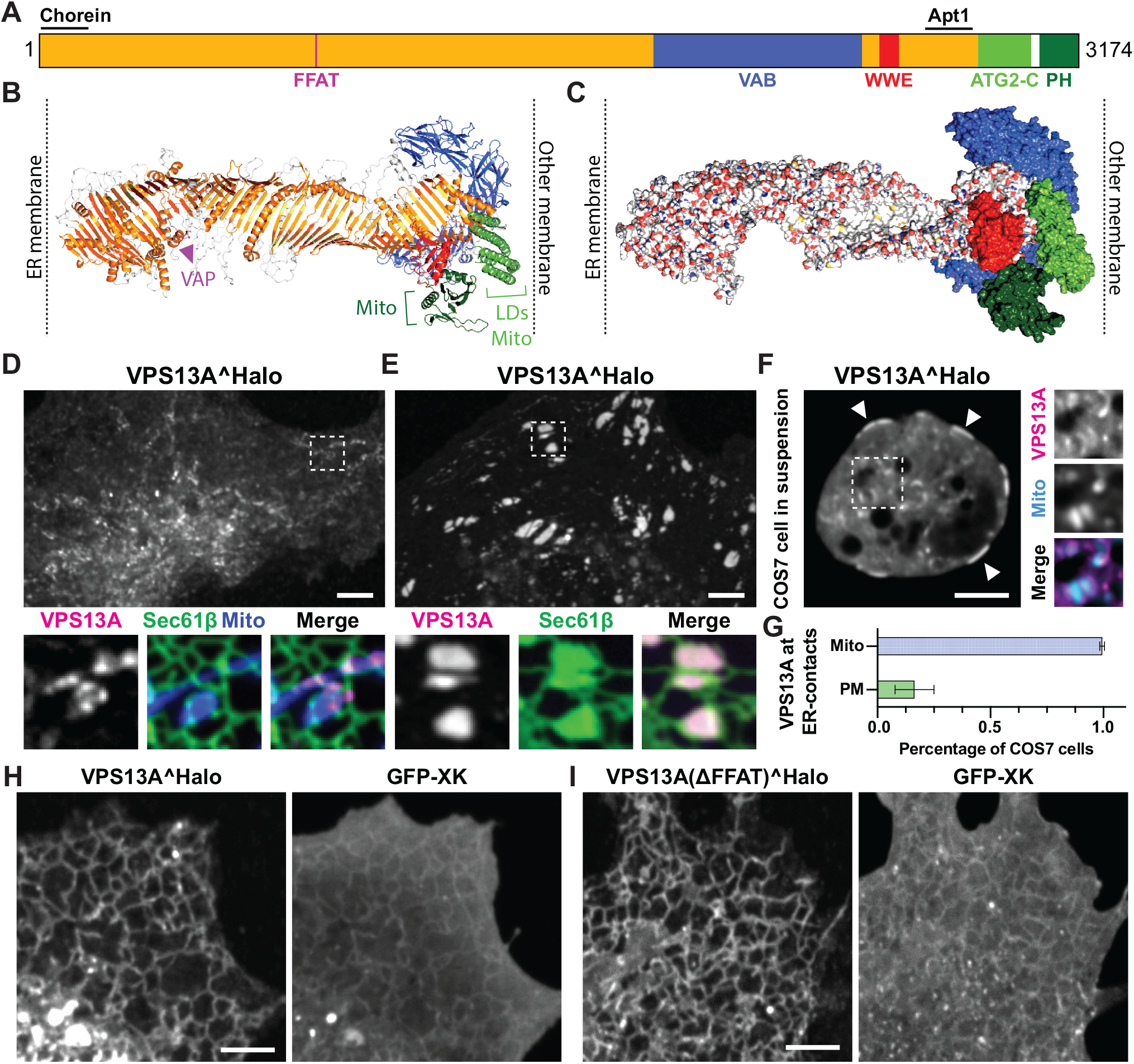
VPS13A is a ≥22 nm long rod that can localize to ER-PM contacts. **(A)** Domain organization of human VPS13A. **(B-C)** Predicted structure of human VPS13A in ribbon (B) and surface (C) representations. In (B), different regions are colored coded as in A and the known binding sites to different organelles or proteins are indicated. LD = lipid droplets. In (C), the *β*-sheet repeat that forms the groove are colored by element: carbon in white, nitrogen in blue (positive charges), oxygen in red (negative charges), and sulphur in yellow, with the white surface representing its hydrophobic floor. Only a portion of such floor is visible due to the twisting of the rod (see also Movie S1). **(D-F)** Confocal images of COS7 cells, attached to the substrate (D and E) or in suspension (F), co-expressing VPS13A^Halo with mito-BFP and/or GFP-Sec61*β* as markers for the mitochondria and ER, respectively. VPS13A localizes to ER-mitochondria contacts (D-F) and, in a small percentage of cells, also to ER-PM (E and F) contacts. The latter can be visualized as patches in proximity to the plasma membrane in the bottom plane of a substrate-attached cell (E) or at the cell cortex in a cell in suspension (F, see arrowheads). Larger magnifications of the areas enclosed by a stippled rectangle in D,E and F are shown at the bottom (D and E) or at the right (F) of the main fields. **(G)** Percentage of COS7 cells expressing VPS13A^Halo that showed a localization to ER-mitochondria and ER-PM contacts. **(H and I)** COS7 cells co-expressing GFP-XK and VPS13A^Halo, with (H) or without (I) its FFAT motif. In these cells the localization of VPS13A to ER-mitochondria contacts is lost and VPS13A is sequestered selectively to the ER, despite XK being present both in the ER (most likely its biosynthetic pool) and in the plasma membrane where it appears as a diffuse fluorescence in these very flat cells. Scale bars=5 μm.

### In a small subset of cells VPS13A localized to ER-PM contacts in a XK-independent way

We and others(9, 14, 16) reported previously that VPS13A binds the outer mitochondrial membrane and lipid droplets via its C-terminal ATG2-C region and PH domains, mediating the formation of VAP-dependent ER-mitochondria and ER-lipid droplets contacts. In agreement with these previous reports, COS7 cells expressing VPS13A^Halo along with Sec61*β*-EGFP and mito-BFP shows a localization of VPS13A at ER-mitochondria contacts (Fig. 1D). Similar results were obtained in RPE1 cells (Fig. S1A). However, we found that in a small population of both COS7 and RPE1 cells VPS13A^Halo is also localized in patches closely apposed to the plasma membrane (PM), in a pattern highly reminiscent of ER-PM contacts sites (Fig. 1D-G and S1B). We hypothesized that XK would be required for this localization. Thus, we used CRISPR-Cas9 in XK loss-of-function studies using RPE1 cells, a model human cell line with a diploid genome. We generated several clones of these cells with different mutations in the XK gene, all leading to early stop codons in the translated polypeptide (Fig. S1C-D). However, a population of cells from these clonal lines still showed VPS13A^Halo patches representative of ER-PM contacts (Fig. S1E-F), suggesting that the protein can bind the PM via an XK-independent mechanism.

### VPS13A binds XK via its PH domain

Next, we performed gain-of-function studies and tested the effect of overexpressing XK on VPS13A^Halo localization in COS7 cells. GFP-XK was localized primarily at the plasma membrane, with a pool being also observed in the ER (probably reflecting its biosynthetic pool) both when expressed alone and when co-expressed with VPS13A^Halo (Fig. 1H). However, in agreement with what had been reported by Park et al. who had shown a relocation of VPS13A from lipid droplets to the ER upon co-expression of XK(9), the localization of VPS13A^Halo was dramatically changed by co-expression with GFP-XK. VPS13A’s presence at mitochondria was completely lost and VPS13A localized instead throughout the ER network where it precisely colocalized with XK(Fig. 1H), surprisingly without an enrichment with XK at the PM. Larger abnormal intracellular ER structures (also referred to as Organized Smooth ER structures (OSERs)(30)) positive for both proteins were additionally observed in some cells, as also reported by Park et al (9) (Fig. S2A and B)(see also below). Importantly, the binding of VPS13A to the ER when co-expressed with XK did not require its binding to VAP, as it was not abolished upon mutation of its VAP-binding FFAT motif (Fig. 1I), suggesting a different mode of binding mediated by a ER-localized pool of XK (see Fig. S2C). Similar results were obtained by co-expressing VPS13A with XKR2, the member of the XK family with the strongest similarity to XK and a reported VPS13A interactor(31), although exogenous VPS13A was still able to localize to ER-PM contacts in a small fraction of XK KO RPE1 cells that had been additionally edited to disrupt the XKR2 gene (Fig S1E and S1G-I). In contrast, expression of other XK proteins (XKR3, XKR6 and XKR8) (Fig. S2D and E) did not affect the localization of VPS13A. Likewise, the localization of other VPS13 proteins tested (VPS13C and VPS13D) was not affected by co-expression of XK (Fig. S2F and G). Collectively, our findings, together with the findings of Park et al.(9), suggest an interaction of VPS13A with XK that competes with the binding to either mitochondria or lipid droplets.

Binding of VPS13A to mitochondria is mediated by both its PH domain and ATG2-C region(14, 16). When expressed alone in either COS7 cells or HEK293 cells, the PH domain of VPS13A (EGFP-PH_VPS13A_) bound mitochondria, but when co-expressed with XK, and surprisingly in contrast to full length VPS13A^Halo, it completely relocalized to XK-containing membranes, both the ER and the PM (Fig. 2A-C). Many PH domains bind directly the lipid bilayer. To confirm that binding of PH_VPS13A_ to the PM in GFP-XK expressing cells reflects binding to XK, an XK construct was generated by inserting in one of its extracellular loops a small sequence (Twin-Strep) recognized by Strep-Tactin XT, a tetrameric protein(32). This XK construct can be used to visualize specifically the plasma membrane (and thus surface-exposed) pool of XK by adding to the medium Strep-Tactin XT conjugated to a fluorophore (Fig. 2D, right panel). Cells expressing such construct together with EGFP-PH_VPS13A_, showed a diffuse distribution of PH_VPS13A_ at the PM. However, addition to the medium of Strep-Tactin XT, resulted in the clustering of XK in parallel with the intracellular co-clustering of EGFP-PH_VPS13A_, proving binding of the two proteins at the PM (Fig. 2D and Movie S2).

**Figure 2.**
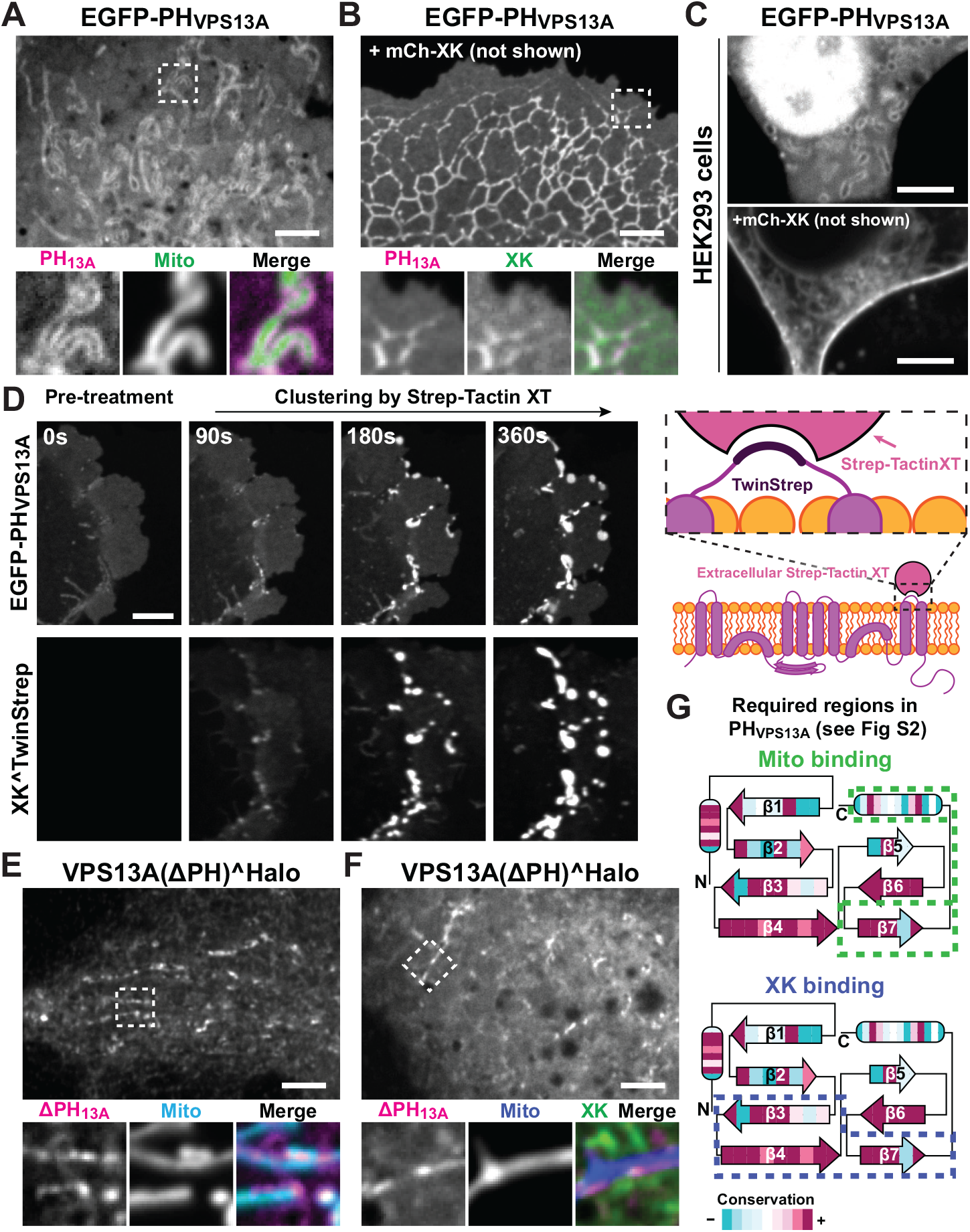
The C-terminal PH domain of VPS13A is necessary and sufficient for the interaction with XK. **(A and B)** Confocal images of COS7 cells showing that EGFP-PH_VPS13A_ binds to mitochondria (A), but relocalizes to ER and PM along with XK when co-expressed with mCh-XK (B). The same is shown in HEK293 cells **(C)**. The diffuse fluorescence of EGFP-PH_VPS13A_ in A represents cytosolic fluorescence as also confirmed by (C) (note that a nuclear fluorescence is often observed with EGFP-tagged PH domains). **(D)** Left: COS-7 cell co-expressing EGFP-PH_VPS13A_ and XK^Twin-Strep showing co-clustering of both proteins upon addition of Strep-Tactin XT conjugated to a DY-649 fluorophore (see also Movie S2). Right: Diagram explaining the binding of XK^Twin-Strep to extracellular Strep-Tactin XT. **(E and F)** COS7 cells co-expressing VPS13A(ΔPH)^Halo with either mito-BFP (E) or Mito-BFP and GFP-XK (F), showing that this construct can bind mitochondria but not XK. **(G)** Cartoon representing the secondary structure of PH_VPS13A_ where the strands and helices (ovals) are colored by conservation among chordates. Stippled lines enclose the portion of the PH domain required for binding to mitochondria or XK. Scale bars=5 μm.

Further supporting the role of the PH domain in the binding to XK, the co-expression of XK did not impact the localization at ER-mitochondria contacts of a VPS13A construct lacking selectively the PH domain (Fig. 2E and F), as the ATG2-C region is sufficient for the binding of VPS13A to the outer mitochondrial membrane. Thus, the PH domain is both necessary and sufficient to recruit VPS13A exclusively to XK-containing membranes. Based on these findings we interpret binding of full length VPS13A to the ER upon expression of XK as due to its PH domain (Fig. 1H) and suggest that the abnormal intracellular ER structures positive for both VPS13A and XK are due to VPS13A-dependent tethering of adjacent ER cisterns (Fig. S2B-C).

The PH domain of VPS13A is highly conserved among chordates, with the highest conservation occurring on one surface (Fig. S3A), suggesting that such surface is the one mediating the competitive interaction with XK and with the yet unknown binding partner on mitochondria. We confirmed this hypothesis by generating chimeras between the PH domains of VPS13A and the one of VPS13C, which does not bind mitochondria or XK (Fig. S3B). These experiments showed that the precise residues involved in the two interactions are not exactly the same, as some constructs were capable of binding XK but not mitochondria (Fig. 2G and S3B-C), and viceversa. However, replacement of the seventh *β*-strand of the PH domain of VPS13A, which lies in the conserved surface, with the corresponding strand of the PH domain of VPS13C, resulted in a cytosolic localization of the domain with or without XK overexpression, suggesting its requirement for the binding to both partners (Fig. 2G and S3B-C).

### Intracellular loops of XK regulate the binding of VPS13A at the PM

We next set out to elucidate how the PH domain of VPS13A binds XK. XK and XKR2 are predicted by AlphaFold2(20, 33) to contain 8 TM helices and 2 hydrophobic helices that are buried in the membrane but are kinked and do not span the entire bilayer (Fig. 3A-B). Such prediction fits with the experimentally determined structures of the two closely related proteins XKR8 and XKR9(34, 35). The main differences between XK/XKR2 and XKR8/XKR9 family members lie in their cytosolically exposed a.a. sequences. Specific features that distinguish XK and XKR2 relative to XKR8 and XKR9 are a *β*-sheet hairpin in the second cytosolic loop and the lack of the caspase-cleavage site in the C-terminal cytosolic tail that is crucial for the activation of the scrambling function of XKR8 and XKR9(6, 10).

**Figure 3.**
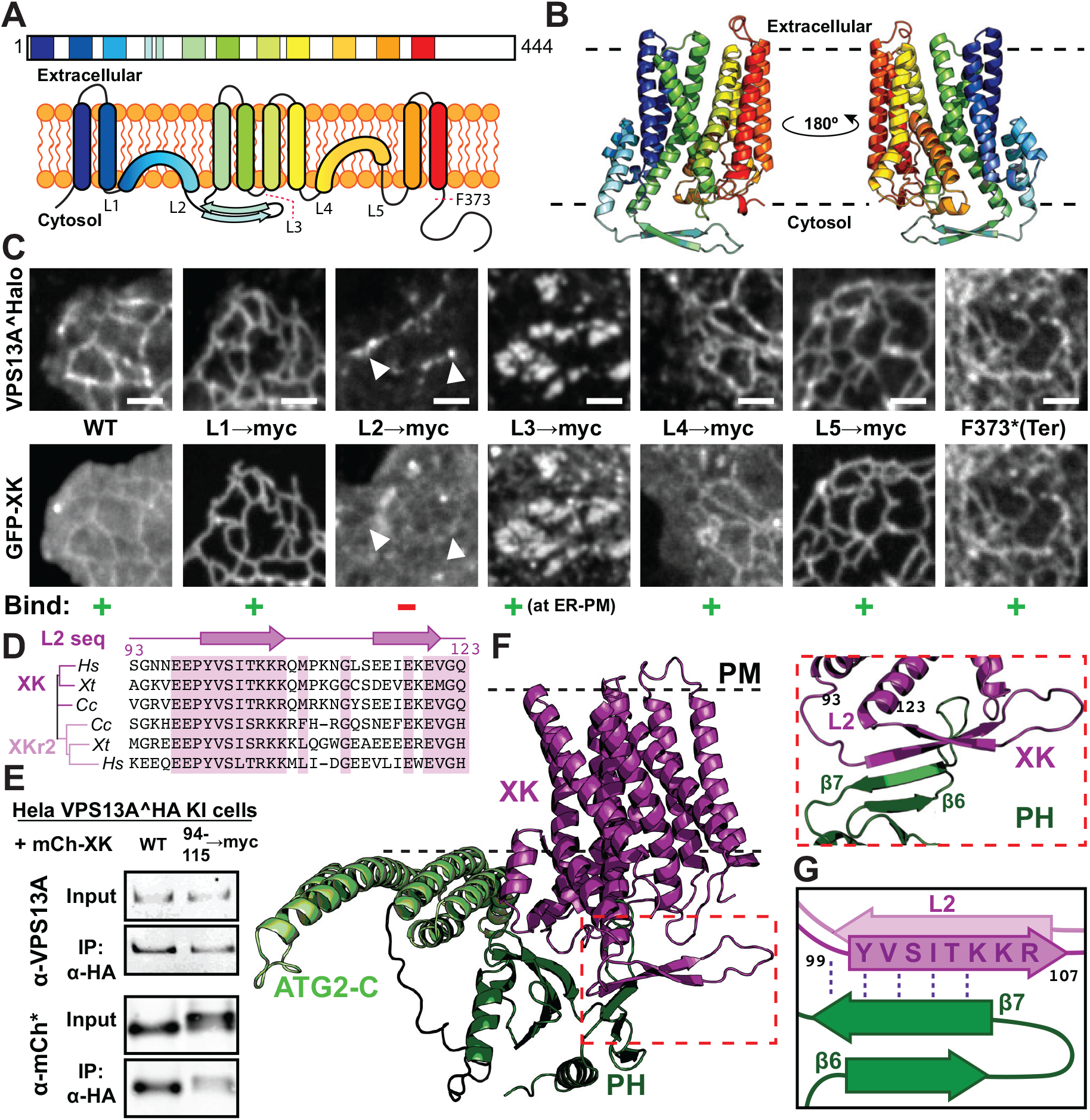
XK binds VPS13A via a conserved cytosolic *β*-hairpin. **(A)** Cartoons representing the transmembrane organization of XK in a linear fashion (top) and in 2D (bottom). In the 2D cartoon, the 5 cytosolic loops of XK are indicated. The position of the phenylalanine (F373) that was mutated to a stop codon in one of the constructs shown in (C) is also indicated. **(B)** Predicted ribbon structure of XK based on AlphaFold2. **(C)** COS7 cells co-expressing VPS13A^Halo and either WT XK or constructs of GFP-XK where the different cytosolic loops have been replaced by the myc tag aminoacid sequence. GFP-XK localizes to both the ER and the PM (which appears as diffuse fluorescence, see legend of Fig. 1H). Mutation of the second cytosolic loop (L2) abolished the interaction of XK with VPS13A (third column). Mutations of L1 and L5 abolished PM labeling suggesting an impact of XK trafficking to the PM, but not the interaction with VPS13A (second and fifth columns). Mutations of L3 led to an accumulation and colocalization of both VPS13A and XK at patches that represent ER-PM contacts (see also Fig 4) Top and bottom images of each column show the same microscopy field. **(D)** Alignment of sequences corresponding to L2 of XK and XKR2 in *Homo sapiens* (*Hs*), *Xenopus tropicalis* (*Xt*) and *Carcharodon carcharias* (*Cc*). The secondary structure of the loop is shown above the sequences, and identical or similar residues are highlighted in purple. **(E)** Immunoprecipitation of VPS13A from the extract of a HeLa cell line where VPS13A was endogenously HA-tagged(14), showing that coprecipitation of mCh-XK is dependent on the N-terminal *β*-strand of L2 of XK. **(F and G)** AlphaFold multimer structural prediction of the interaction between XK and the ATG2-C-PH domains of VPS13A. The N-terminal *β*-strand of L2 of XK is predicted to interact with the seventh *β*-strand of the PH, in agreement with experimental data. Scale bars=2 μm.

To identify the region responsible for binding VPS13A, we replaced, individually, the cytosolic loops of XK with the myc-tag a.a sequence (Fig. 3C). This analysis revealed that the second cytosolic loop conserved in XK and XKR2 (Fig. 3D), i.e. the *β*-sheet hairpin (Fig. 3D), was crucial for the interaction between the two proteins, as expression of an XK construct in which this loop was replaced with a myc tag (EGFP-XK_L2→myc_) did not compete with the localization of co-transfected VPS13A to ER-mitochondria contacts (Fig. 3C, third column). The importance of the second loop for the interaction was supported by anti-HA immunoprecipitation from knock-in HeLa cells expressing VPS13A tagged with 2 HA epitopes at the endogenous locus(14) and co-expressing either wild type XK or XK with a myc tag replacing the N-terminal portion of the hairpin in this loop (mCh-XK_94-115→myc)_ (Fig. 3E). Predictions using Alphafold-Multimer(36) also indicated that the binding of PH_VPS13A_ to XK is mediated at least in part by residues in the seventh *β* strands of the PH domain and in the N-terminal *β*-strand of the second cytosolic loop of XK (Fig. 3F and G), in full agreement with our experimental data. Moreover, in the predicted XK-VPS13A complex the a-helices of the ATG2-C domain are oriented with the hydrophobic surface facing the XK-containing membrane (Fig. S3D and E), raising the possibility that ATG2-C could partially penetrate the bilayer and thus synergize with the XK-PH_VPS13A_ interaction in in the membrane recruitment of VPS13A.

AlphaFold2 predictions of XK alone(20, 33) also showed an intramolecular interaction between the *β*-strand of the second cytosolic loop of XK that binds PHVPS13A and the third cytosolic loop (Fig. 4A). We speculated that such interaction could potentially regulate the accessibility of the second cytosolic loop to VPS13A. We tested this hypothesis by co-expressing VPS13A^Halo with XK constructs where the residues implicated in this intramolecular binding were mutated - either in the second (KKR→AAA) or third (EYE→AAA) cytosolic loop. Strikingly in these cells we found a massive accumulation of VPS13A^Halo at patches with the localization and morphology of ER-PM contacts, where the two proteins colocalized (Fig. 4B-D). These patches were confirmed to represent ER-PM contacts as when the XK_KKR→AAA_ was additionally modified by addition of the Twin-Strep tag for recognition by extracellular Strep-Tactin XT and co-expressed with full length VPS13A and the ER protein VAP, all three proteins precisely colocalized (Fig. 4E-F). The colocalization of the two proteins at ER-PM contacts was in striking contrast to the colocalization of the two proteins selectively in the ER when VPS13A^Halo was co-expressed with WT XK. Collectively, these findings confirmed that VPS13A is capable of tethering the ER to PM via binding to XK, but suggested a yet unknown regulatory mechanism by which this binding is regulated.

**Figure 4.**
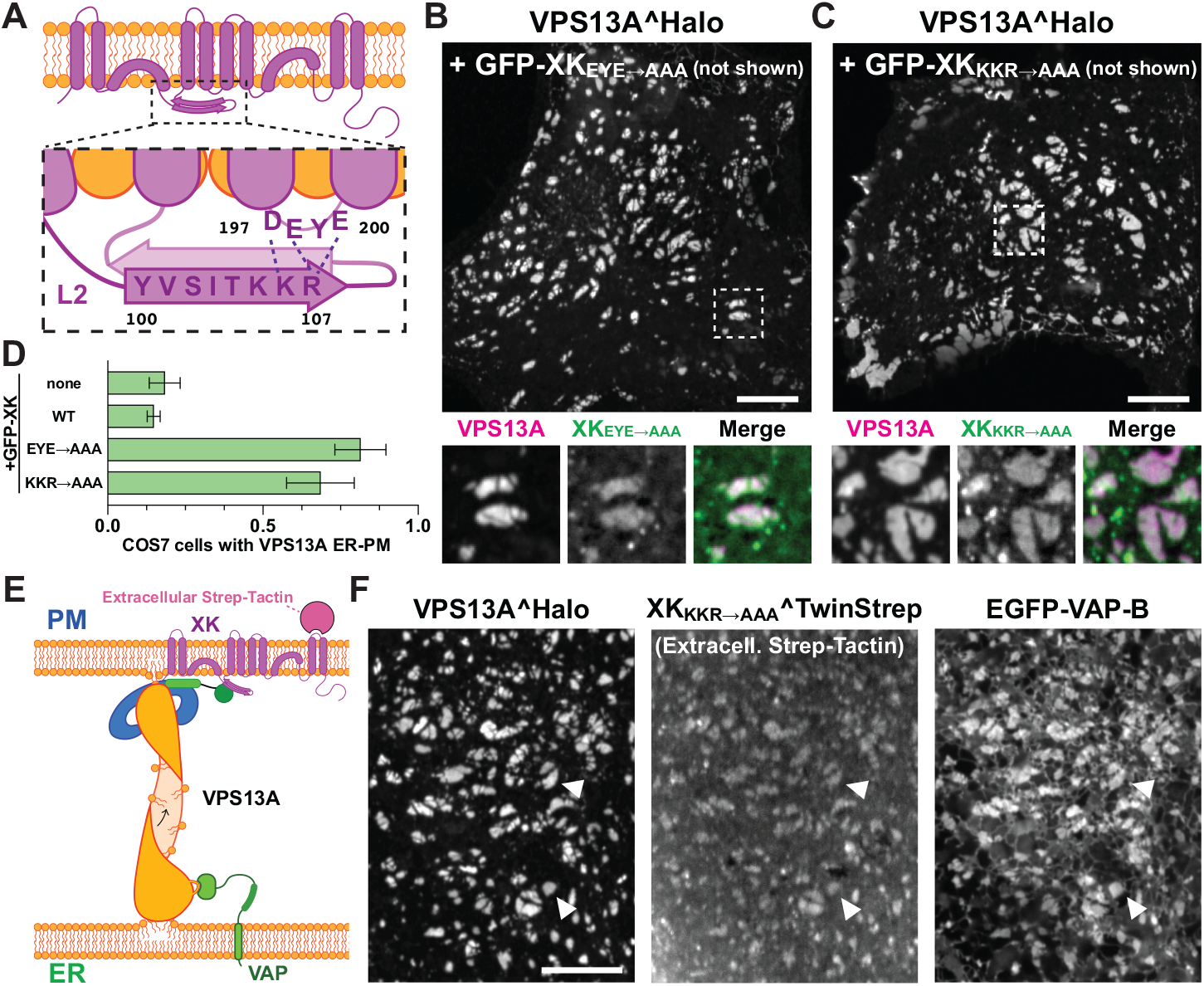
An intramolecular interaction between two cytosolic loops of XK impacts the binding of VPS13A to XK at the PM. **(A)** Cartoon representation of XK structure showing residues in the N-terminal *β*-strand of L2 and in L3 that are predicted to interact based on AlphaFold predictions. **(B and C)** COS7 cells co-expressing VPS13A^Halo and constructs of GFP-XK where the a.a E198, Y199 and E200 in L3 (B) or K105, K106 and R107 in L2 (C) were replaced with alanines, showing presence of VPS13A at ER-PM contacts. **(D)** Percentage of COS7 cells not expressing XK or expressing WT or mutant XK that shows VPS13A at ER-PM contacts. **(E)** Cartoon of the ER-PM tether mediated by VPS13A via the interaction with VAP in the ER and XK in the PM. **(F)** COS7 cells co-expressing VPS13A^Halo, XK_KKR→AAA_^Twin-Strep (visualized by extracellular Strep-Tactin XT), and EGFP-VAP-B, showing that the three proteins colocalize at ER-PM contacts. Scale bars=10 μm.

### Implications for the pathogenesis of chorea-acanthocytosis and McLeod syndrome

The PH domain of VPS13A is of crucial importance for ChAc pathogenesis, as at least 9 (out of more than 50 reported(37)) patient mutations were reported to occur in this domain, which is encoded by the last three exons of the VPS13A gene. Interestingly, one of several VPS13A splice variants lacks the PH domain as it uses an alternative exon 69 (out of 72 in total) that encodes a short disordered region rich in acidic residues followed by a stop codon (Fig. 5A). When expressed in COS7 cells, this splice variant (isoform B) did not bind XK but was still localized at ER-mitochondria contacts (Fig. 5B), similarly to the artificially engineered VPS13A construct lacking the PH domain (Fig. 2F and G). Intriguingly, previous studies reported this splice isoform to be brain specific and to be the predominant VPS13A splice variants expressed in mouse brain tissue(38), including the striatum where neurodegeneration is observed in neuroacanthocytosis patients. As this finding questioned a role of the VPS13A-XK interaction in the neurological manifestations of ChAc and MLS, we investigated expression of the PH domain containing isoform A (the longer isoform) in brain tissue. Using qPCR we found that in the human caudate the longer isoform A is expressed at higher levels than the shorter isoform (Fig. 5C)

**Figure 5.**
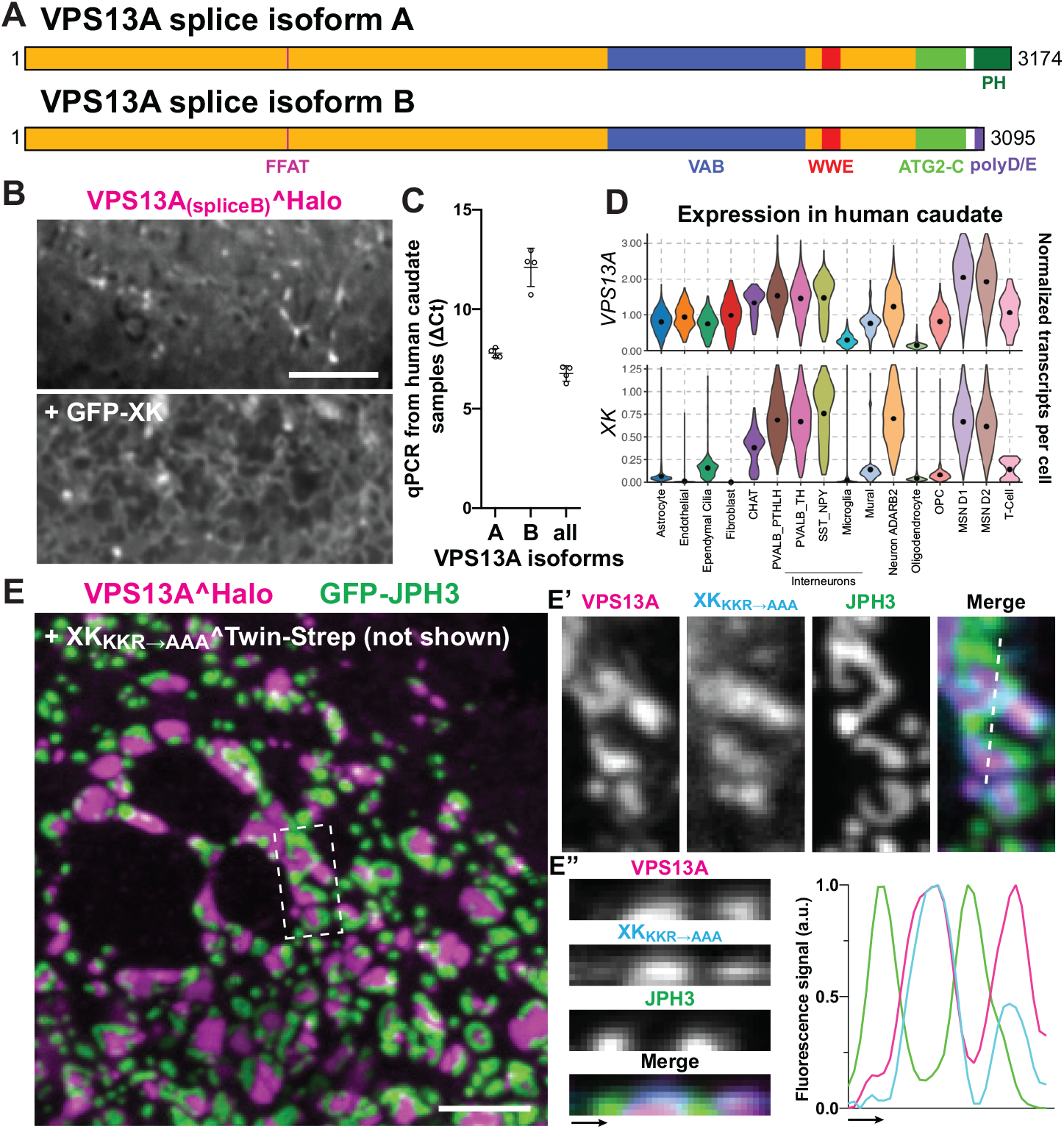
VPS13A and XK are expressed in human striatum. **(A)** Domain organization of two splice variants of VPS13A. **(B)** COS7 cells co-expressing VPS13A_(splice variant B)_^Halo and GFP-XK, showing no colocalization as expected due to the lack of the PH domain. The top and bottom images shows the same microscopy field. **(C)** qPCR quantitation of transcript levels of splice variants A and B of *VPS13A* in human caudate samples. Splice variant A is expressed at higher levels than variant B, as opposed to what was reported in mice. *VPS13A* isoform levels in each case were normalized to housekeeping gene *ACTB*, with ΔCt for each isoform calculated from the qPCR cycle threshold (Ct) of the *VPS13A* isoform versus the *ACTB* Ct. **(D)** Violin plots of single nucleus RNA-seq data of human caudate samples showing preferential expression of *VPS13A* and *XK* in neurons, including Medium Spiny Neurons (MSNs) where *VPS13A* has the highest expression levels. **(E)** COS7 cells co-expressing XK_KKR→AAA_^Twin-Strep, VPS13A^Halo and GFP-JPH3. The figure, which shows a view in a plane close to the substrate, reveals that both VPS13A^Halo and GFP-JPH3 localize to ER-PM contacts but do not overlap. E’ shows at high magnification the area enclosed by a rectangle in E and E” shows an orthogonal view along the dashed line of E’. Scale bars=5 μm.

We next examined expression of VPS13A and XK in different cell populations of the striatum by analyzing previously obtained single nuclei RNAseq data(39). We found that the expression of VPS13A and XK are highly correlated in human striatum and that D1 and D2 medium spiny neurons (MSNs) of this region, i.e. the neuronal population that degenerates in ChAc and MLS, are among the striatal cells with the highest expression of both genes (Fig. 5D). Interestingly, the same co-enrichment of VPS13A and XK in medium spiny neurons does not occur in mice striatum (Fig. S4A). This, in addition to the higher enrichment of the shorter variant in mice striatum(38), may explain the lack of major striatal neurodegeneration in VPS13A loss-of-function mice models(40, 41)

Mutation of two other genes result in syndromes (collectively referred to as neuroacanthocytosis syndromes) similar to ChAc and MLS. These genes are *PANK2*, which encodes a mitochondrial protein whose mutations results in Panthothenate Kinase-Associated Neurodegeneration (PKAN)(42), and *JPH3*, which encodes junctophilin 3, an ER to PM tether whose mutations result in a Huntington Disease Like condition (HDL2)(43). In the human caudate *PANK2* is ubiquituously expressed, with enrichment in non-neuronal cells, whereas expression of *JPH3* is restricted to neurons, including medium spiny neurons (Fig. S4B). The *JPH3* gene encodes two junctophilin 3 isoforms. Interestingly, the main isoform is a tail-anchored transmembrane protein of the ER that binds acidic lipids in the PM via N-terminally localized MORN motifs and thus is localized at ER-PM contacts(44, 45). The other iso-form is a much shorter protein of unknown function that comprises a small portion of the N-terminal region. The *JPH3* mutations responsible for HDL2 consist of repeat expansions in an exon used selectively in the shorter isoform and it was reported that this expansion partially impairs the function of the main isoform(46). Thus, a potential functional relationship of junctophilin 3 with the tether mediated by VPS13A and XK is plausible. In COS7 cells, both VPS13A^Halo and GFP-JPH3 localized to patches representing ER-PM contacts, although the patches positive for VPS13A were adjacent but not precisely overlapping with those positive for Junctophilin (Fig. 5E). A similar segregation of a VPS13 family member (VPS13C) and another membrane contact site protein (PDZD8) was observed at ER-endolysosome contacts, possibly reflecting, at least in part, size exclusion due to the length of VPS13 proteins(23). VPS13A is ∼22 nm long, whereas contacts formed by junctophilins in muscle cells has been reported to be around 12nm(47). In spite of the lack of precise colocalization, the independent binding of junctophilin 3 to the PM may facilitate the encounter of VPS13A with XK and thus the formation and/or stabilization of XK-VPS13A

## Discussion

Our study proves that VPS13A can interact with XK at the plasma membrane thus strengthening the hypothesis that these two proteins are functional partners. It also suggests that such interaction, which we show to be mediated by the PH domain of VPS13A and the second cytosolic loop of XK, may be subject to regulation. Previously, we and other had found that VPS13A localizes at ER mitochondria contacts and, when lipid droplets are present, also at ER-lipid droplets contacts(14, 16). The localization with XK at the plasma membrane is an additional localization that does not exclude, but compete with, these other interactions. The occurrence of multiple sites of action for VPS13A, one of the four VPS13 paralogues expressed by mammalian cells, is reminiscent of the properties of the single yeast VPS13 which functions at multiple sites.

XK was discovered as a binding partner to the Kell glycoprotein, an antigen in the membrane of erythrocytes(48). Its function, however, remained elusive for many years until XKr8, one of nine XK paralogs, was reported to be necessary for the apoptosis-dependent scrambling of lipids in the plasma membrane, a process that results in the surface exposure of PtdSer(49). Surface exposed PtdSer, in turn, acts as the “eat me signal” which is read by surrounding cells to promote the engulfment and degradation of the dying cell(7, 49). Following this initial study, other XK paralogs, XKR4 and XKR9 were also shown to scramble PM lipids in response to apoptotic signals(6). The structures of XKR8 and XKR9 were recently solved by Cryo-EM(34, 35). AlphaFold2-based prediction(20, 33) of the structure of XK suggests an overall very similar fold (10)and thus a similar scramblase function, as confirmed by recent in vitro studies(13). However, XK differs from XKR8 and XKR9 in the cytosolic loops and C-terminal tail. In XKR8 and XKR9, a helix in the C-terminal tail folds under the transmembrane core, potentially stabilizing it in an inactive conformation(34, 35). During apoptosis, the tail is cleaved by caspase-3 resulting in a conformational change of the protein that allows phospholipid scrambling between the two bilayer leaflets. Neither the tail, nor the cleavage site, are conserved in XK or XKR2. However, the second cytosolic loop, which is the VPS13A-binding *β*-strand hairpin unique to XK and XKR2, is predicted by Alphafold2 to form a direct intramolecular interaction with the third cytosolic loop, possibly playing a similar role to the C-terminal tail of XKR8 in stabilizing an inactive conformation. In fact, the amino acids responsible for this intramolecular interaction are precisely the same ones whose mutations cause the recruitment of full length VPS13A to XK at the plasma membrane. Our findings open the possibility that binding of VPS13A to XK may regulate its activity by inducing a conformational change similar to the one caused by caspase cleavage of the C-terminal tail of XKR8 and XKR9. A potentially similar scenario occurs with XKR4, which requires the binding of a caspase-cleaved peptide of the nuclear protein XRCC4 to its cytosolic loops to activate scrambling(50).

It remains unclear why PH_VPS13A_ binds WT XK irrespective of its location, both at the PM and in the ER (the biosynthetic pool of XK), while exogenous full length VPS13A binds XK in the ER, but not in the plasma membrane, unless the intramolecular interaction within the cytosolic domain of XK is disrupted by mutations. Clearly, the functional partnership of XK and VPS13A implies their interaction at plasma membrane in WT cells. Elucidating the mechanisms underlying this unexpected finding, which suggests a regulation of the intramolecular interaction of XK relevant for the binding to full length VPS13A, but not of its PH domain alone, remains a priority for the future. Another open question is what recruits exogenous VPS13A to the plasma membrane in a small subset of cells without overexpression of XK and even in XK KO or XK/XKR2 double KO cells. Such recruitment points to the existence of another (likely regulated) binding site for VPS13A at the plasma membrane.

A potential scenario is that VPS13A delivers phospholipids from the ER to the cytosolic leaflet of the PM and that the activity of XK allows equilibration of the abundance of phospholipids in the two leaflets. This equilibration would result in surface exposure of PtdSer if not efficiently compensated by the action of flippases. This scenario is supported by a recent study identifying VPS13A and XK as genes essential for the ATP-dependent exposure of PtdSer in T cells expressing the purinergic receptor P2×7(10). Studies in model organisms are in agreement with the hypothesis that XK and a VPS13 family protein plays a role in PtdSer externalization. *Drosophila* flies lacking Vps13, which is the ortholog of both VPS13A and VPS13C in this organism, have defects in the removal of nurse cell corpses in late stage egg chambers(51). Interestingly, an accumulation of Vps13 was observed close to the plasma membrane of nurse cells, where cisterns likely to represent cortical ER (and thus sites of ER-PM apposition) are also present. Such cisterns are absent in Vps13-/- nurse cells which fail to be engulfed by neighboring follicle cells after their dumping of cytosol to the oocyte has concluded(51). Although it has not been established that these nurse cells expose PtdSer, the externalization of this phospholipid is the signal that triggers engulfment by surrounding cells throughout eukaryotes(7). Moreover, follicle cells require Draper, a surface receptor for PtdSer, to engulf apoptotic cells(52, 53). Similarly, persistent cell corpses are reported as a phenotype in the *C. elegans* mutant (T08G11.1 is the homolog of VPS13A and VPS13C)(54).

Given the clinical similarities of ChAc and MLS patients, it is likely that abnormal lipid dynamics by the VPS13A-XK partnership lies at the core of the pathogenesis of both diseases. The co-expression of both proteins in medium spiny neurons of the caudate nucleus, i.e. the brain cells that degenerate in both conditions, is consistent with this possibility. Interestingly, *JPH3*, another neuroacanthocytosis gene also expressed in these neurons, encodes for an ER-PM tether, suggesting a potential functional relationship with the VPS13A-XK complex at these sites. However, how a defect in lipid scrambling in the plasma membrane of striatal neurons may be involved in neurodegeneration remains to be investigated. One possibility is that the absence of VPS13A or XK may impair normal homeostasis of the of the PM, due to either a defect in lipid delivery or of a coupling between lipid delivery and lipid scrambling to equilibrate bilayer leaflets. Another possibility is that neurodegeneration may be due to a defect in PtdSer externalization. Externalization of this lipid by neuronal cells was shown to be implicated in removal of dead cells and in synaptic pruning, suggesting a physiological significance of this process(55, 56). The impairment of PtdSer exposure may prevent normal clearance of cell debris that need to be eliminated. Future studies should address a potential role of VPS13A and XK in PtdSer exposure in human medium spiny neurons, where the expression of the two proteins is enriched and highly correlated.

In the context of neurodegeneration, it is of interest that a WWE domain is present within the structure of VPS13A, as well as of VPS13C(23), whose mutations also results in neurodegeneration (Parkinson’s disease)(57). WWE domains were shown to bind Poly-ADP-Ribose (PAR)(28), a molecule implicated in glutamate excitotoxicity-mediated neuronal death (also termed PARthanathos, due to its dependence on PAR)(58). Upon excessive stimulation of NMDA-receptors, DNA damage activates the protein PARP-1, which synthesizes PAR. Excess PAR, in turn, translocates from the nucleus to the cytosol, where it triggers the release of apoptotic-inducible factor (AIF) from mitochondria(59). Cultured neurons exposed to excess glutamate have been shown to expose PtdSer via a still unclear caspase-independent process(60) and several neurodegeneration conditions are thought to be at least partially caused by glutamate excitotoxicity(61).

A major open question concerns the role of the partnership between XK and VPS13A in the hematological manifestations of ChAc and MLS (abnormally shaped erythrocytes), as mature erythrocytes do not contain ER and thus ER-PM junctions. It remains possible that the defect may occur during erythrocytes maturation in erythroblastic islands, where erythroblasts expel their nuclei surrounded with a layer of plasma membrane, which then externalize PtdSer and are engulfed by macrophages(62). However, like XK, which together with Kell is part of the Kx blood antigen, VPS13A is present in the plasma membrane of adult erythrocytes. As these cells do not contain ER, the significance of such a presence remains unclear.

## Methods

### DNA plasmids

The following constructs were generated in our lab: VPS13A^Halo (Addgene #118759), VPS13C^Halo, VPS13D^Halo (Addgene #174108), EGFP-E-Syt1(Addgene #66830), EGFP-JPH3 and EGFP-VAP-B. GFP-XK, GFP-XKR2 and XKR8-myc were obtained from Genscript. cDNA sequences encoding XKR3 and XKR6 were purchased from Genscript and Origene, respectively, and cloned to a pEGFP-C1 (Addgene) backbone. All other constructs were generated from the above mentioned constructs using In-Fusion cloning (Takara) or site-directed mutagenesis (QuikChange II XL; Agilent technologies). The primers and/or restriction enzymes are described in the supplementary table. Mito-BFP was a gift from G. Voeltz (Addgene #49151).

### Antibodies and reagents

Primary antibodies used were anti-VPS13A (NBP1-85641; Novus Biological) and anti-RFP (600-401-379; Rockland Inc). Halo tag ligands were a kind gift from L. Lavis (Janelia Research Campus, Ashburn, VA). The Twin-Strep tag and Strep-Tactin XT protein fluorescently conjugated with DY-649 were purchased from IBA Lifesciences.

### Immunoprecipitation of VPS13A

VPS13A^2xHA KI HeLa cells (previously generated in our lab(14) were transiently transfected using FuGene HD (Promega) with either mCh-XK or mCh-XK_94-115→myc_. After 48 hours, cells were lysed in a buffer containing 1%Triton X-100 for 30min, the lysate was further diluted to a concentration of 0.33%Triton X-100 and incubated with anti-HA magnetic beads (Pierce) for 2h. Beads were washed and the samples were eluted with SDS-sample buffer.

### Cell culture and transfection

COS7, HEK293, RPE1(CRL-4000) and HeLa cells were obtained from ATCC. Cells were cultured at 37ºC and 5% CO2 in DMEM (or DMEM:F12 in the case of RPE1) containing 10%FBS, 1mM sodium pyruvate, 100U/ml penicillin, 100mg/mL streptomycin and 2mM L-glutamine (all from Gibco). RPE1 cells For imaging experiments, cells were seeded on glass-bottomed dishes (MatTek) at a concentration of 50-75×103 cells per dish, transiently transfected using FuGene HD (Promega) and imaged ≥48 hours later.

### Microscopy

#### Live cell imaging

Just before imaging, the growth medium was removed and replaced with pre-warmed Live Cell Imaging solution (Life technologies). Imaging was carried out at 37ºC and 5%CO2. Spinning-disk confocal microscopy was performed using an Andor Dragonfly system equipped with a PlanApo objective (63X, 1.4NA, Oil) and a Zyla sCMOS camera. For obtaining images of COS7 cells in suspension, COS7 were trypsinized and re-plated in MatTeks 10min before imaging. *Immunofluorescence*. Cells were fixed with 4% PFA, permeabilized with 0.1% Triton-X-100 and blocked with 5% BSA. Primary (anti-c-Myc, 9E10; Santa Cruz) and secondary (Goat anti-Mouse IgG, Invitrogen) antibodies were used at 1:200 and 1:1000 dilutions, respectively, in buffer containing 0.1% Triton-X-100 and 5% BSA.

### XK KO cell line generation

sgRNAs targeting the human XK gene were generated using the IDT-DNA online tool. RPE1 cells were transiently transfected using FuGene HD (Promega) with plasmids containing Cas9 and the sgRNAs (the backbone used was plasmid PX459, a gift from Feng Zhang (Addgene #62988)). Transfected cells were then selected after 24h with 2ug/ml Puromycin. The selection antibiotic was removed after 72 hours, and clonal cell populations were isolated approximately a week after. Mutations in the XK gene were confirmed by PCR and sequencing using the primers listed in the supplementary table.

### qPCR from human caudate samples

Human caudate analyses were conducted as exempt human research, as this was secondary research using biospecimens not specifically collected for this study. Four pathologically normal samples were obtained from the NIH NeuroBioBank using appropriate de-identification and under consent. Tissue was disrupted for RNA isolation using the TissueLyser (QIAGEN, Hilden, Germany) for 2 × 2 min at 20 Hz as recommended by the manufacturer and bulk RNA extracted using the RNeasy Lipid Tissue Mini Kit (QIAGEN, Hilden, Germany). For qRT-PCR, TaqMan probes and Universal Master Mix were used (ThermoScientific) and run on a StepOnePlus system (ThermoScientific, Rockford, IL). The TaqMan probes used for specific detection of splice isoforms were: for A: Hs00364243_m1, for B: Hs00362923_m1, or detection of both isoforms: Hs00362891_m1. All experiments were run in technical triplicates for each biological sample and normalized to *ACTB*, with Ct for each isoform calculated from the Qpcr cycle threshold (Ct) of the *VPS13A* isoform versus the *ACTB* Ct.

## Acknowledgements

We thank L. Zhang, M. Hanna and J.H. Park for discussion. The manuscript file uploaded to biorXiv was generated using the LaTeX template adapted by S. Royle available at https://github.com/quantixed/manuscript-templates. Work in the P.D.C. lab was supported in part by NIH grants NS36251, DA018343 and DK45735, the Parkinson Foundation (PF-RCE-1946) and by grant #2020-221912 from the Chan Zuckerberg initiative DAF, an advised fund of Silicon Valley Community Foundation. Work in the M.H. lab was supported by NIH grant R01NS100802.

## Author contributions

AGS and PDC conceptualized the project. AGS, YW, SSP, FJG, JNE, ML and BU designed and performed experiments. AGS, YW, SSP, FJG, MK, MH and PDC analysed data. AGS and PDC wrote the original draft of the manuscript which was then reviewed and edited by all authors.

## Supplementary

**Figure S1.**
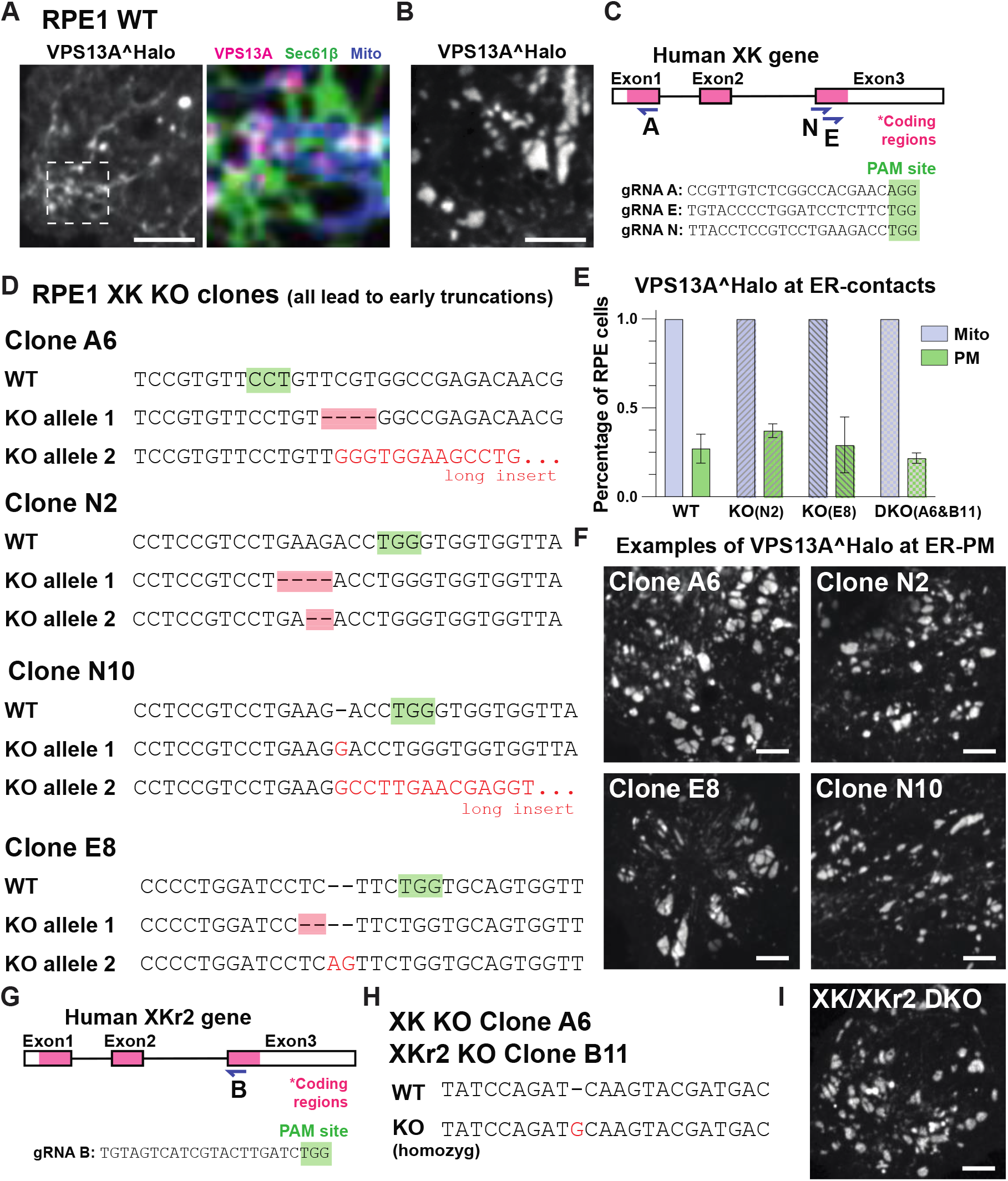
VPS13A can localize to ER-PM contacts in XK KO cells. (A-B) Confocal images of RPE1 cells, expressing VPS13A^Halo showing localization to ER-mitochondria (A) or ER-PM contacts (B). (C) Schematic representation of the XK gene. Three gRNA sequences that target different exons were used. (D) Four different XK KO clonal populations were selected and tested. (E) Percentage of RPE cells expressing VPS13A^Halo that showed a localization to ER-mitochondria and to ER-PM contacts. (F) Representative examples of XK KO RPE cells expressing VPS13A^Halo and showing a localization to ER-PM. (G) Schematic representation of the XKr2 gene. (H) Sequence of mutation introduced in XKr2 gene in XK KO cells (clone A6) background. (I) Representative example of XK/XKr2 DKO cell expressing VPS13A^Halo and showing a localization to ER-PM. Scale bars=5μm.

**Figure S2.**
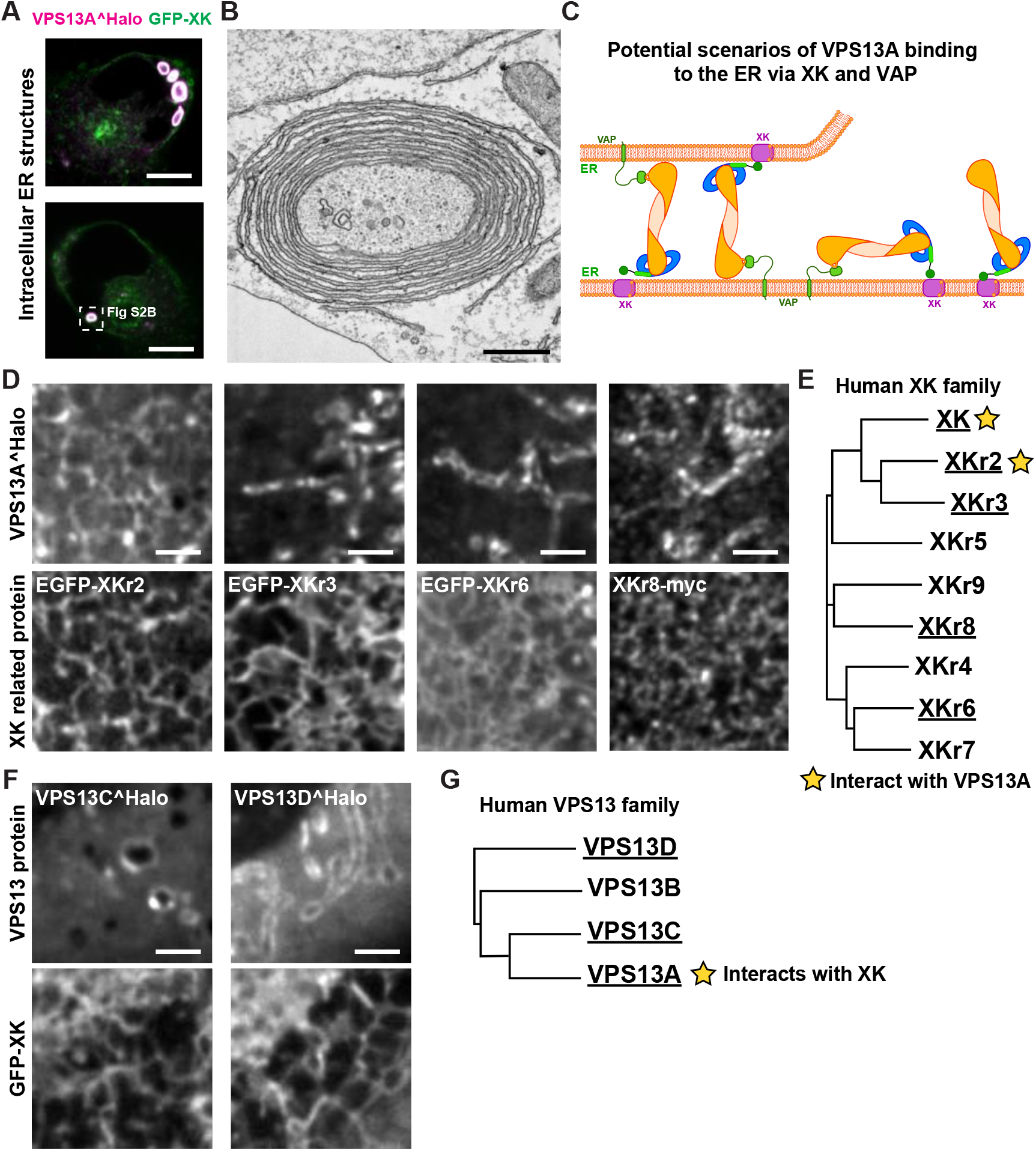
VPS13A interacts with XK and XKr2. (A-C) Binding of VPS13A to XK in the ER. (A) Confocal images of COS7 cells with high enrichment of co-expressed VPS13A^Halo and GFP-XK in round and oval hollow structures that Correlative-Light Electron Microscopy (CLEM) revealed to be circular stack of ER cisternae (B). We speculate that these structures, which resemble OSERs (Organized Smooth ER)(30)) represent appositions of ER membranes generated as shown by the cartoon shown in (C). Note in (C) that the binding of VPS13A does not require VAP. (D) COS7 cells coexpressing VPS13A^Halo (top images) with XKR2 and other proteins of the XK-related family (bottom images). Top and bottom images shows the same microscopy field. VPS13A^Halo is recruited to the ER by XKr2 but not by XKr3, XKr6 or XKr8. (E) Human XK family tree. The underlined members were tested for interaction as shown in (D). (F) COS7 cells co-expressing GFP-XK with other VPS13 family members, showing that XK does not interfere with VPS13C’s localization to endolysosomes or VPS13D’s to mitochondria. Top and bottom images shows the same microscopy field (G) Human VPS13 family tree. Scale bars= 10μm for A, 2.5μm for D and F.

**Figure S3.**
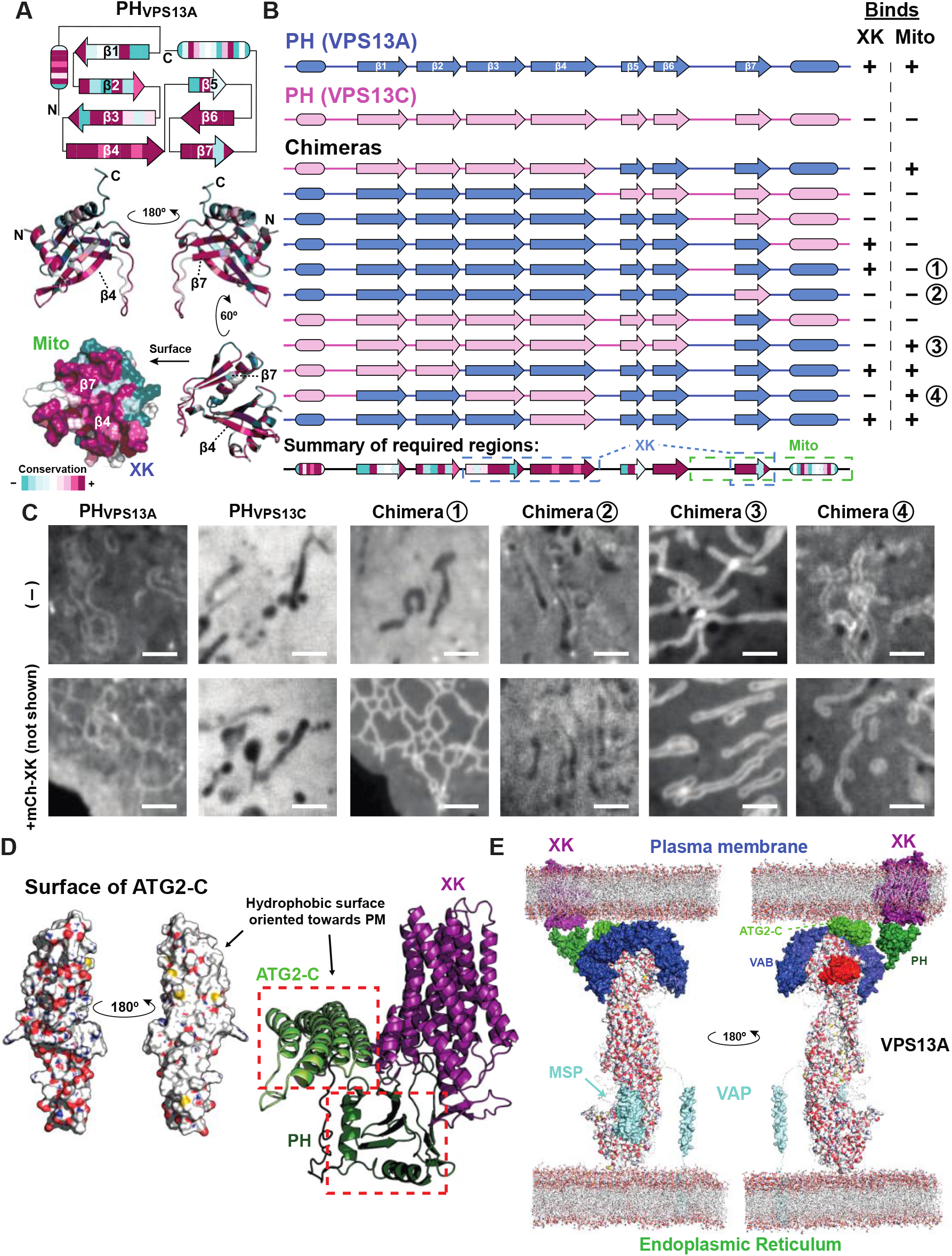
The PH domain of VPS13A interacts with XK and mitochondria via a conserved surface. (A) Cartoon representing the secondary structure of PHVPS13A (top) and ribbon and surface representations of its tertiary structures predicted by Alphafold (bottom), all colored by conservation among chordates. The surface representation indicates the conserved surface that binds both XK and mitochondria, although by only partially overlapping regions. (B) Diagram indicating all the chimeras between the PH domains of VPS13A and VPS13C that we have tested, and their binding to XK or to mitochondria. (C) COS7 cells expressing the indicated PH domain construct by itself (top row) or with mCh-XK co-expression (bottom row). Scale bar=2.5μm. (D) Alphafold multimer structural prediction of the interaction between XK and the ATG2-C-PH fragment of VPS13A, as shown in Fig. 3F. The ATG2-C domain has a highly hydrophobic surface which is oriented towards the XK-containing membrane. (E) Schematic of the predicted organization of VPS13A at the ER-PM interface via the interaction with the MSP domain of VAP and with XK. Only one monomer of VAP (a dimeric protein) is shown.

**Figure S4.**
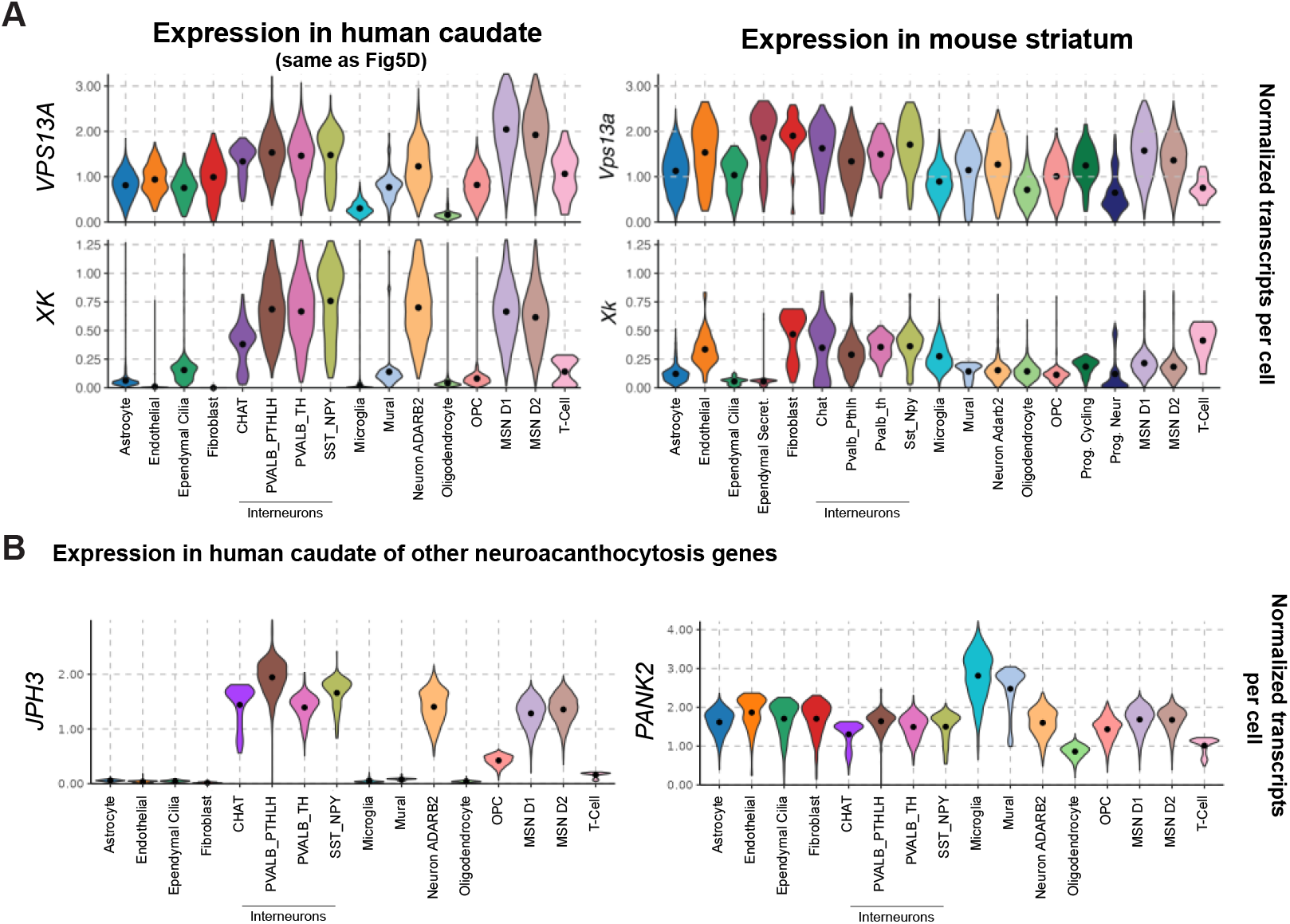
Expression of neuroacanthocytosis genes in human caudate. (A) Expression levels of *VPS13A* and *XK*, as revealed by snRNA-seq, in human (left, same as Fig 5D) and mice (right) striatal cells. *VPS13A* and *XK* are enriched in neurons in human samples whereas their expression is more uniform across cell types in mice. (B) Expression pattern in the human caudate of the two other genes also associated with neuroacanthocytosis syndromes, *JPH3* and *PANK2*. Expression of *JPH3* is restricted to neurons whereas that of *PANK2* is ubiquitous and higher in non-neuronal cell populations.

**Table S1.**
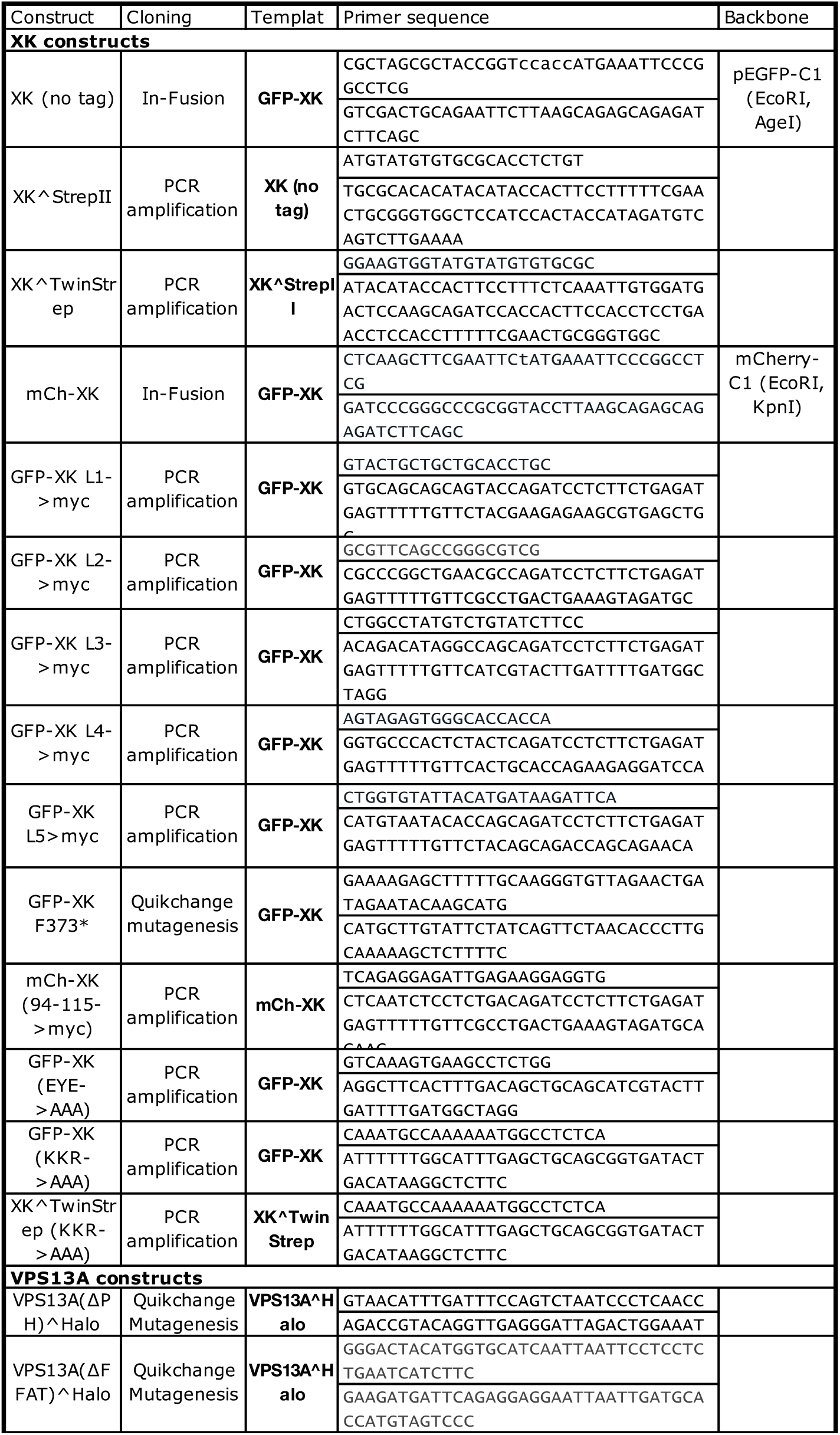
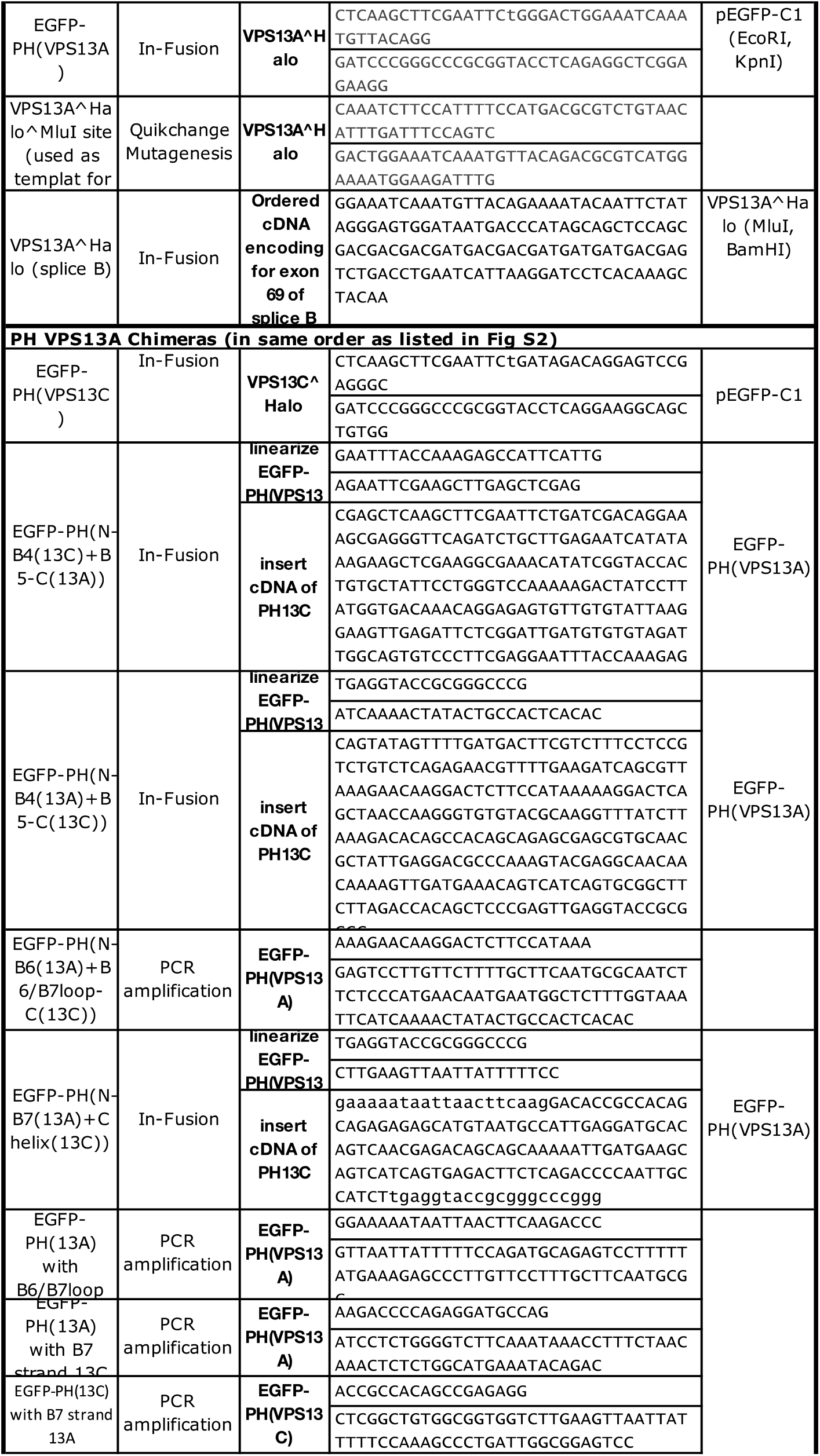
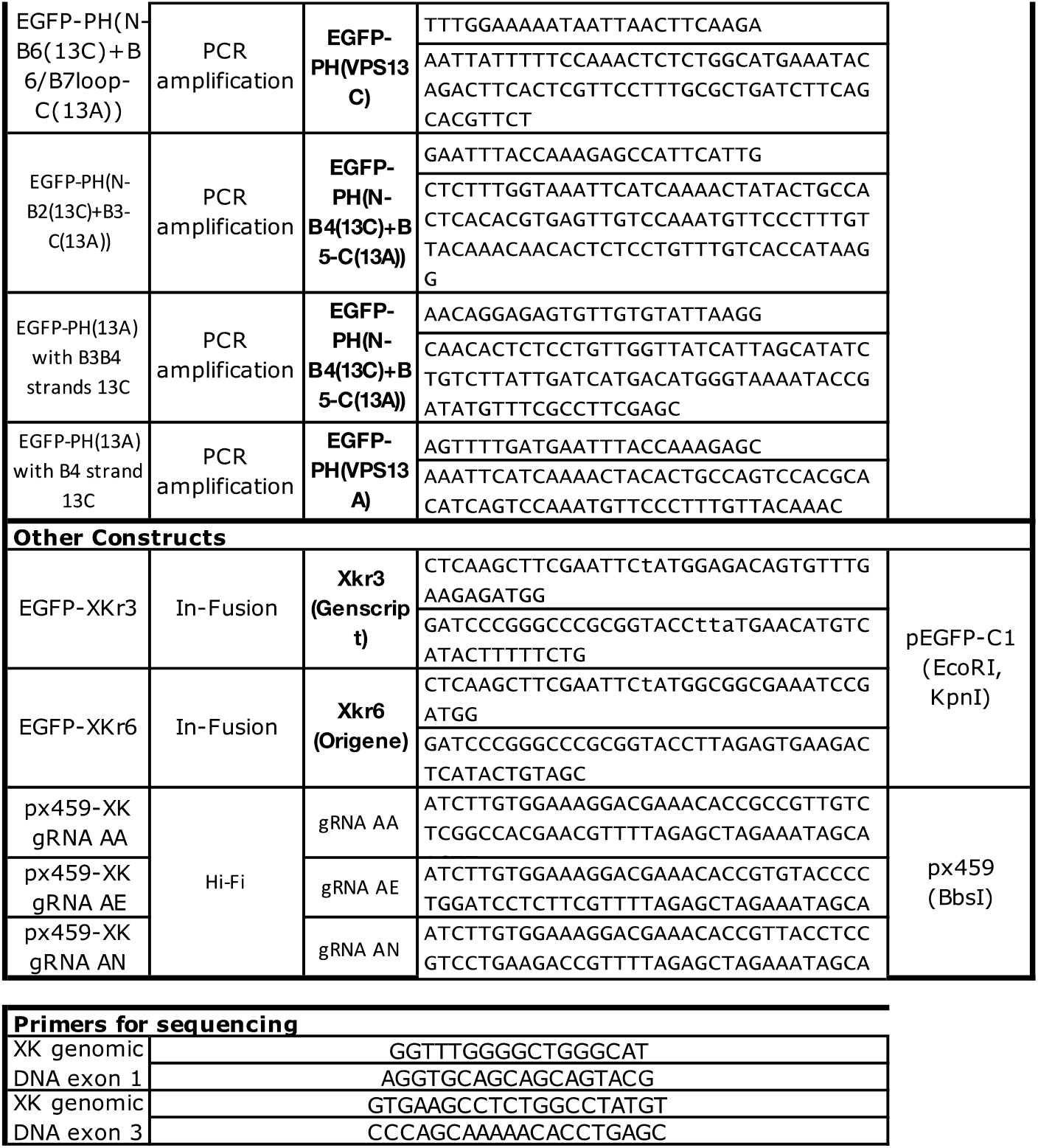
Oligonucleotides used in this study.

### Supplementary Movies (mp4 files uploaded separatly)

**Movie S1. VPS13A is a** ∼**22 nm long rod with a hydrophobic cavity**. Rotating VPS13A structure in ribbon and surface representation. The ribbon representation is colored as in Figure 1A and the surface representation by element: carbon in white, nitrogen in blue (positive charges) and oxygen in red (negative charges).

**Movie S2. The PH domain of VPS13A interacts with XK at the plasma membrane**. COS-7 cell co-expressing EGFP-PHVPS13A and XK^Twin-Strep showing co-clustering of both proteins upon addition of Strep-Tactin XT conjugated to a DY-649 fluorophore at time=0s. Scale bar=5μm.

## References

1. L. Rampoldi, et al., A conserved sorting-associated protein is mutant in chorea-acanthocytosis. Nat. Genet. 28, 119–120 (2001).

2. S. Ueno, et al., The gene encoding a newly discovered protein, chorein, is mutated in chorea-acanthocytosis. Nat. Genet. 28, 121–122 (2001).

3. S. K. Dziurdzik, E. Conibear, The Vps13 Family of Lipid Transporters and Its Role at Membrane Contact Sites. Int J Mol Sci 22, 2905 (2021).

4. M. Leonzino, K. M. Reinisch, P. D. Camilli, Insights into VPS13 properties and function reveal a new mechanism of eukaryotic lipid transport. Biochimica Et Biophysica Acta Mol Cell Biology Lipids 1866, 159003–159003 (2021).

5. M. Ho, et al., Isolation of the gene for McLeod syndrome that encodes a novel membrane transport protein. Cell 77, 869–880 (1994).

6. J. Suzuki, E. Imanishi, S. Nagata, Exposure of phosphatidylserine by Xk-related protein family members during apoptosis. J Biological Chem 289, 30257–30267 (2014).

7. S. Nagata, J. Suzuki, K. Segawa, T. Fujii, Exposure of phosphatidylserine on the cell surface. Nature Publishing Group 23, 952–961 (2016).

8. Y. Urata, et al., Novel pathogenic XK mutations in McLeod syndrome and interaction between XK protein and chorein. Neurology Genetics 5, e328 (2019).

9. J.-S. Park, A. M. Neiman, XK is a partner for VPS13A: a molecular link between Chorea-Acanthocytosis and McLeod Syndrome. Molecular Biology of the Cell 31, 2425–2436 (2020).

10. Y. Ryoden, K. Segawa, S. Nagata, Requirement of Xk and Vps13a for the P2×7-mediated phospholipid scrambling and cell lysis in mouse T cells. Proc National Acad Sci 119, e2119286119 (2022).

11. N. N. Noda, Atg2 and Atg9: Intermembrane and interleaflet lipid transporters driving autophagy. Biochimica Et Biophysica Acta Bba - Mol Cell Biology Lipids 1866, 158956 (2021).

12. A. Ghanbarpour, D. P. Valverde, T. J. Melia, K. M. Reinisch, A model for a partnership of lipid transfer proteins and scramblases in membrane expansion and organelle biogenesis. Proc National Acad Sci 118, e2101562118 (2021).

13. J. Adlakha, Z. Hong, P. Li, K. M. Reinisch, Structural and biochemical insights into lipid transport by VPS13 proteins. Biorxiv, 2022.03.11.484024 (2022).

14. N. Kumar, et al., VPS13A and VPS13C are lipid transport proteins differentially localized at ER contact sites. J Cell Biol 217, 3625–3639 (2018).

15. S. Muñoz-Braceras, A.R. Tornero-Écija, O. Vincent, R. Escalante, VPS13A is closely associated with mitochondria and is required for efficient lysosomal degradation. Dis Model Mech 12, dmm036681 (2019).

16. W. M. Yeshaw, et al., Human VPS13A is associated with multiple organelles and influences mitochondrial morphology and lipid droplet motility. eLife 8, 728 (2019).

17. X. Liu, et al., An AP-MS- and BioID-compatible MAC-tag enables comprehensive mapping of protein interactions and subcellular localizations. Nature Communications 9, 1188 (2018).

18. C. D. Go, et al., A proximity-dependent biotinylation map of a human cell. Nature 595, 120–124 (2021).

19. A. Guillén-Samander, et al., VPS13D bridges the ER to mitochondria and peroxisomes via Miro. J Cell Biol 220, e202010004 (2021).

20. J. Jumper, et al., Highly accurate protein structure prediction with AlphaFold. Nature 596, 583–589 (2021).

21. M. De, et al., The Vps13p-Cdc31p complex is directly required for TGN late endosome transport and TGN homotypic fusion. J Cell Biol 216, 425–439 (2017).

22. P. Li, J. A. Lees, C. P. Lusk, K. M. Reinisch, Cryo-EM reconstruction of a VPS13 fragment reveals a long groove to channel lipids between membranes. J Cell Biol 219, 3593 (2020).

23. S. Cai, et al., In situ architecture of the lipid transport protein VPS13C at ER-lysosomes membrane contacts. Biorxiv, 2022.03.08.482579 (2022).

24. T. J. Melia, K. M. Reinisch, A possible role for VPS13-family proteins in bulk lipid transfer, membrane expansion and organelle biogenesis. J Cell Sci 135 (2022).

25. S. E. Murphy, T. P. Levine, VAP, a Versatile Access Point for the Endoplasmic Reticulum: Review and analysis of FFAT-like motifs in the VAPome. Biochim. Biophys. Acta 1861, 952–961 (2016).

26. B. D. M. Bean, et al., Competitive organelle-specific adaptors recruit Vps13 to membrane contact sites. J Cell Biol 217, 3593–3607 (2018).

27. D. R. Fidler, et al., Using HHsearch to tackle proteins of unknown function: A pilot study with PH domains. Traffic 17, 1214–1226 (2016).

28. L. Aravind, The WWE domain: a common interaction module in protein ubiquitination and ADP ribosylation. Trends Biochem Sci 26, 273–275 (2001).

29. W. Rzepnikowska, et al., Amino acid substitution equivalent to human chorea-acanthocytosis I2771R in yeast Vps13 protein affects its binding to phosphatidylinositol 3-phosphate. Hum Mol Genet 26, ddx054 (2017).

30. E. L. Snapp, et al., Formation of stacked ER cisternae by low affinity protein interactions. J Cell Biol 163, 257–269 (2003).

31. E. L. Huttlin, et al., Dual proteome-scale networks reveal cell-specific remodeling of the human interactome. Cell 184, 3022-3040.e28 (2021).

32. T. G. M. Schmidt, et al., Development of the Twin-Strep-tag® and its application for purification of recombinant proteins from cell culture supernatants. Protein Expres Purif 92, 54–61 (2013).

33. M. Varadi, et al., AlphaFold Protein Structure Database: massively expanding the structural coverage of protein-sequence space with high-accuracy models. Nucleic Acids Res 50, D439–D444 (2022).

34. T. Sakuragi, et al., The tertiary structure of the human Xkr8–Basigin complex that scrambles phospholipids at plasma membranes. Nat Struct Mol Biol 28, 825–834 (2021).

35. M. S. Straub, C. Alvadia, M. Sawicka, R. Dutzler, Cryo-EM structures of the caspase-activated protein XKR9 involved in apoptotic lipid scrambling. Elife 10, e69800 (2021).

36. R. Evans, et al., Protein complex prediction with AlphaFold-Multimer. Biorxiv, 2021.10.04.463034 (2022).

37. C. Dobson-Stone, et al., Mutational spectrum of the CHAC gene in patients with chorea-acanthocytosis. Eur J Hum Genet 10, 773–781 (2002).

38. E. Mizuno, et al., Brain-specific transcript variants of 5 and 3 ends of mouse VPS13A and VPS13C. Biochem Bioph Res Co 353, 902–907 (2007).

39. H. Lee, et al., Cell Type-Specific Transcriptomics Reveals that Mutant Huntingtin Leads to Mitochondrial RNA Release and Neuronal Innate Immune Activation. Neuron 107, 891-908.e8 (2020).

40. H. Sakimoto, M. Nakamura, O. Nagata, I. Yokoyama, A. Sano, Phenotypic abnormalities in a chorea-acanthocytosis mouse model are modulated by strain background. Biochem Bioph Res Co 472, 118–24 (2016).

41. K. Peikert, et al., Therapeutic targeting of Lyn kinase to treat chorea-acanthocytosis. Acta Neuropathologica Commun 9, 81 (2021).

42. S. J. Hayflick, Unraveling the Hallervorden-Spatz syndrome: pantothenate kinase-associated neurodegeneration is the name. Curr Opin Pediatr 15, 572–577 (2003).

43. S. E. Holmes, et al., A repeat expansion in the gene encoding junctophilin-3 is associated with Huntington disease–like 2. Nat Genet 29, 377–378 (2001).

44. H. Takeshima, S. Komazaki, M. Nishi, M. Iino, K. Kangawa, Junctophilins A Novel Family of Junctional Membrane Complex Proteins. Mol Cell 6, 11–22 (2000).

45. M. Nishi, H. Sakagami, S. Komazaki, H. Kondo, H. Takeshima, Coexpression of junctophilin type 3 and type 4 in brain. Mol Brain Res 118, 102–110 (2003).

46. A. I. Seixas, et al., Loss of junctophilin-3 contributes to huntington disease-like 2 pathogenesis. Ann Neurol 71, 245–257 (2012).

47. H. Takeshima, Intracellular Ca2+ store in embryonic cardiac myocytes. Front Biosci 7, d1642 (2002).

48. C. M. Redman, et al., Biochemical studies on McLeod phenotype red cells and isolation of Kx antigen. Brit J Haematol 68, 131–136 (1988).

49. J. Suzuki, D. P. Denning, E. Imanishi, H. R. Horvitz, S. Nagata, Xk-related protein 8 and CED-8 promote phosphatidylserine exposure in apoptotic cells. Sci New York N Y 341, 403–6 (2013).

50. M. Maruoka, et al., Caspase cleavage releases a nuclear protein fragment that stimulates phospholipid scrambling at the plasma membrane. Mol Cell 81, 1397-1410.e9 (2021).

51. A. I. E. Faber, et al., Vps13 is required for timely removal of nurse cell corpses. Dev Camb Engl 147, dev191759 (2020).

52. J. I. Etchegaray, et al., Draper acts through the JNK pathway to control synchronous engulfment of dying germline cells by follicular epithelial cells. Development 139, 4029–4039 (2012).

53. T. T. Tung, et al., Phosphatidylserine recognition and induction of apoptotic cell clearance by Drosophila engulfment receptor Draper. J Biochem 153, 483–91 (2013).

54. WormBase version WS283. Available on: http://www.wormbase.org/db/get?name=WBGene00011629;class=Gene (Accessed on March 21, 2022).

55. E. C. Damisah, et al., Astrocytes and microglia play orchestrated roles and respect phagocytic territories during neuronal corpse removal in vivo. Sci Adv 6, eaba3239 (2020).

56. N. Scott-Hewitt, et al., Local externalization of phosphatidylserine mediates developmental synaptic pruning by microglia. Embo J 39, e105380 (2020).

57. S. Lesage, et al., Loss of VPS13C Function in Autosomal-Recessive Parkinsonism Causes Mitochondrial Dysfunction and Increases PINK1/Parkin-Dependent Mitophagy. Am. J. Hum. Genet. 98, 500–513 (2016).

58. K. K. David, S. A. Andrabi, T. M. Dawson, V. L. Dawson, Parthanatos, a messenger of death. Frontiers Biosci Landmark Ed 14, 1116–28 (2009).

59. H. C. Kang, T. M. Dawson, V. L. Dawson, “Excitotoxic Programmed Cell Death Involves Caspase-Independent Mechanisms” in Acute Neuronal Injury, D. G. Fujikawa, Ed. (Springer, 2018), pp. 79–88.

60. H. Wang, et al., Apoptosis-Inducing Factor Substitutes for Caspase Executioners in NMDA-Triggered Excitotoxic Neuronal Death. J Neurosci 24, 10963–10973 (2004).

61. X. Dong, Y. Wang, Z. Qin, Molecular mechanisms of excitotoxicity and their relevance to pathogenesis of neurodegenerative diseases. Acta Pharmacol Sin 30, 379–387 (2009).

62. H. Yoshida, et al., Phosphatidylserine-dependent engulfment by macrophages of nuclei from erythroid precursor cells. Nature 437, 754–758 (2005).

